# Passage through submersible and Archimedes pumps results in decreased migration survival in coho salmon yearlings

**DOI:** 10.64898/2026.09.14.751498

**Authors:** Stephanie Lingard, Hollis Kinnard, Andrew Lotto, Kamil Schzlata, Erika Eliason, Enda Murphy, Dan Straker, Eduardo Martins, Scott Hinch

## Abstract

Flood-control pumpstations compromise connectivity and survival for migratory fish globally, yet field evaluations of modern “fish-friendly” designs remain scarce. We evaluated out-migration survival and behavior of yearling coho salmon (*Oncorhynchus kisutch*) entrained through an open-Archimedes screw station (Hammersley) and a conventional radial station (Barrowtown) in the lower Fraser River floodplain, British Columbia. Using acoustic telemetry over two migration seasons (2024–2025), we tracked smolts across an 84-km corridor to New Westminster and modeled apparent survival using a Cormack–Jolly–Seber framework. Baseline natural mortality in downstream controls averaged 60% across years (cumulative survival: ~40%). Entrainment through the conventional radial station caused 87.5% mortality. The open-Archimedes station conferred an improvement over the conventional facility but still induced 11.5% mortality. Notably, pump-induced losses at the Archimedes pump were predominantly latent: while acute survival through immediate tailrace receivers was high (95–98%), survival deficits diverged downstream in the mainstem Fraser River. Entrainment also prolonged out-migration, causing up to a 10-day delay in 2024 driven by forebay holding and depressed downstream transit speeds. Our findings demonstrate that while open-Archimedes screws improve survival relative to legacy radial impellers, they still impose substantial latent mortality. In heavily altered basins where baseline smolt mortality is already high, additive losses from drainage infrastructure may exacerbate survival bottlenecks.

## Introduction

Flood management is critical to public safety and agriculture, but traditional flood control systems compromise the productivity of aquatic ecosystems and fisheries resources. Pumpstations are deployed throughout floodplains globally to discharge tributary waters across dikes into major downstream waterbodies. In many regions, pumpstations are reaching end-of-life and require urgent modernisation. The demand for pumping capacity is rapidly increasing as a result of climate-induced shifts in hydrology and flood risk (Nawarat et al., 2026; Zhu et al., 2026). However, entrainment through these stations causes substantial injury and mortality to migratory fish (Bierschenk et al., 2019; Bolland et al., 2019; Bruneel et al., 2024), including Pacific salmon (*Oncorhynchus* spp.). Despite growing awareness of this conflict among managers, empirical data necessary for the design of biologically safe pumpstations remain scarce.

Pumpstations installed prior to modern environmental management frameworks, hereafter referred to as “conventional”, employ submersible axial or radial flow pump with tight clearances and high rotational speeds that inflict significant damage to entrained fish (Buysse et al., 2014). Fish passing through pumpstations encounter both mechanical and hydraulic forces that have the potential to cause immediate (direct) mortality through major injury processes such as amputation, organ damage, and barotrauma (Cox et al., 2023). Hydraulic structures also often inflict injuries that increase susceptibility to additional selection pressures (e.g., predation [Mesa, 1994]) and ultimately result in latent mortality (Arkoosh et al., 2006; Budy et al., 2002). The potential impacts of hydraulic infrastructure on fish are reviewed in detail in Cox et al. (2023). To address these issues, manufacturers have created devices with modified impellers, lower rotational speeds, chambers that direct fish away from the impeller, and unpressurized screw systems that eliminate the risk of barotrauma (Abdelghafar et al., 2026; Bierschenk et al., 2019; Van Esch, 2012). These modified devices are marketed as fish-friendly, however, the term has no standardized definition in most jurisdictions. Safety testing of fish-friendly pumps is typically completed by manufacturers on a limited range of species, life stages and design configurations (Bolland & Franklin, 2025). Responses to mechanical and hydrological forces are diverse among species (Bruneel et al., 2024; Helfrich et al., 2001; Pauwels et al., 2020) and stations designs are often bespoke to a specific site. The development of fish safe pumps and infrastructure design guidelines has been impeded by both these intricacies as well as the paucity of literature on this subject.

Archimedes Screw pumps are often promoted as the most fish-friendly design because they operate at constant atmospheric pressure which eliminates the risk of barotrauma. Screw pumps are available in several designs: (1) open-screw where a screw rotates in a concrete housing; (2) closed-unfixed where the screw rotates in a metal tube; (3) closed-fixed where the housing and the screw are welded together and rotate as one unit. Salmon passing through a station with a closed-fixed screw experienced mostly dermal injuries and high immediate survival (> 99%; McNabb et al., 2003). The two unfixed Archimedes designs have not been tested for salmon safety, but tests of an open-screw with other teleost species produced mortality rates of 19-37% (Pauwels et al., 2020). Dermal injuries (scale loss, operculum damage, fin tears, contusions) were most commonly reported in both McNabb et al. (2003) and Pauwles et al. (2020) suggesting the main mechanisms of injury are turbulence, shear force, and collisions (Cox et al., 2023). Fish passing conventional stations experience similar forces with the additional stress of rapid pressure fluctuations that cause barotrauma (Yang et al., 2021). In contrast to the 1-37% mortality estimated for Archimedes screw pumps, grey literature indicates salmon entrained in conventional stations in British Columbia experience > 80% immediate mortality (Thomson, Alan, 1999).

In British Columbia, open Archimedes Screw pumps are commonly installed when municipalities are seeking to improve survival outcomes for Pacific Salmon. Over 100 flood pumps are operated in the floodplain of the lower Fraser River (Finn et al., 2021). In this region, pumps are operated daily to manage tidal flux, or seasonally to manage run-off. Seasonal pumps are typically operated during the spring freshet and fall rainy season. Juvenile salmon emigrating to marine waters are often exposed to pumps as they leave tributaries of the Fraser River. Much of the pump infrastructure in the Fraser River requires urgent upgrades or an expansion of pump facilities to meet climate driven shifts in sea level, hydrology, and irrigation demand (Ali, A & Tanveer, S, 2026; Stakiw et al., 2026). Pacific salmon are also experiencing large scale declines due to the cumulative effects of habitat loss (Finn et al., 2021) and environmental stressors (Walsh et al., 2020).

Latent mortality is an often-overlooked component of infrastructure risk to fish populations, but its quantification is integral to effective conservation actions (Sandford et al., 2012). Two previous studies of salmon survival at pumpstations attempted to quantify latent mortality through 96-hr holding studies (Helfrich et al., 2001; McNabb et al., 2003). However, fish in holding pens are not exposed to predators and challenging environmental or hydrological conditions while recovering from a passage event, as is common in the wild (Mensinger et al., 2024). Acoustic telemetry is a useful tool for quantification of latent mortality following exposures to anthropogenic and environmental stressors (Burnett et al., 2017; Hinch et al., 2025). Here we apply acoustic telemetry to evaluate survival in juvenile coho salmon (*O. kisutch*) following entrainment through an open Archimedes screw pump and an adjacent conventional radial pumpstation. By comparing survival of fish entrained through each pump station to a downstream control group, we disentangle natural mortality from pump-induced mortality in a heavily altered floodplain.

## Methods

### Site description

#### Hammersley

The Hammersley pump station regulates water levels in Mountain Slough, Hogg Slough, and several drainage ditches in the District of Kent (Figures 1, S1). The waterways are part of the historical floodplain of the Fraser River that are now contained by a diked drainage system to facilitate agricultural and urban development. The pumpstation provides flood protection for over 300 km^2^ of developed land. In 2019, the station was upgraded from pressurized impeller pumps to two Landustrie Archimedes Screw pumps (LANDY-270/122/1402-A4), each with a nominal discharge capacity of 2.5 m3/s. Under normal freshet conditions, the station operates with only one pump at a time while both pumps operate simultaneously during higher-flow flood conditions. Each Archimedes screw sits in a cast-in-place concrete trough, with a trash rack at the screw inlet. Water is conveyed through the rotating screw and discharged over a 3.04 m-high sill into a downstream concrete discharge basin. From the basin, water is conveyed through an outlet pipe to the downstream drainage system. The outlet pipe has a slope of approximately 4.5 degrees from the horizontal and is protected by a trash rack at its downstream end. The two pumps produce distinct hydraulic conditions within the discharge basin before their flows converge in the shared outlet pipe. Water discharged from one pump flows directly into the outlet pipe (outflow_straight), while water from the second pump, is directed into an end wall which creates a zone of turbulence and vorticity before turning the corner and draining through the outlet pipe (outflow_corner). The pumpstation operates when the Fraser River at Hope (Water Survey of Canada Gauge: 08MF005) discharge exceeds 2800 m^3^s^−1^. The Hammersley pumpstation also includes two sluice gates (3 m in diameter) that allow the stream to discharge through the dike when flood risk is low (discharge < 2800 m^3^s^−1^). The period of the year the pump is operated, generally, extends from early May to late August.

**Figure 1.**
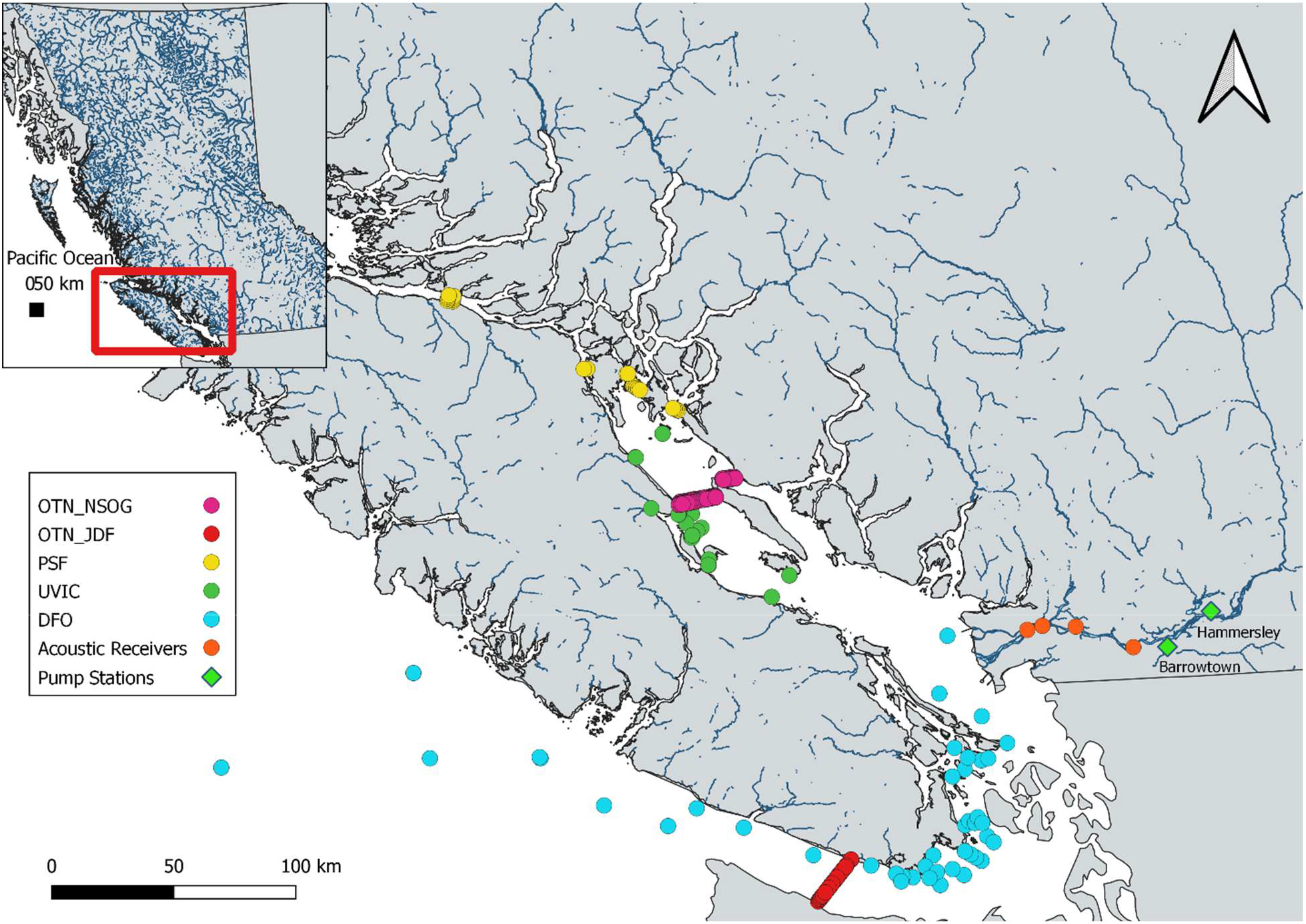
Map of study area in southern British Columbia. Diamonds indicate locations of the two pumpstations evaluated; circles indicate locations of acoustic receivers. Colours pertain to the various organizations to which receivers belong to. OTN=Ocean Tracking Network, PSF=Pacific Salmon Foundation, UVIC=University of Victoria, DFO=Fisheries and Oceans Canada. Orange receivers were operated by Kintama Research Ltd on behalf of the study authors.

#### Barrowtown

The Barrowtown pumpstation is one of the largest in North America. The station was constructed in the 1920’s to prevent flooding in the Sumas Prairie region of Abbotsford, Canada (Figure 1). The station was upgraded in the 1980s to have four mixed-flow pumps manufactured by Stork-Pompen. The pumps operate at either high (117 RMP; 360 m^3^m^−1^) or slow (176 RPM; 600 m^3^m^−1^) speeds. The pumpstation has complex operations and manages the elevation of the Sumas River and Sumas Canal (Figure S2). Water can be pumped from either of these water bodies into the lower unregulated portion of the Sumas River which drains into the Fraser River ~ 4.5 km downstream of the pumpstation. During freshet, when juvenile salmon are migrating to the ocean, the predominate operations are from the Sumas Canal into the Fraser River. Fish enter the system through a screened intake and are transported via suction through piping before passing through the pump. After passing through the body of the pump, fish are then directed to the outlet through another section of piping. The outlet is a flap gate that is submerged at 2.5 m depth during spring freshet.

### Fish tagging

Juvenile coho salmon were obtained from the Chehalis River Hatchery and implanted with acoustic transmitters (V6-69 kHz Innovasea Systems Inc, 6.3 × 13 mm, 0.5 g in air) in May of 2024 (*n*=198) and 2025 (*n*=429; Table 1). A minimum mass of 8.5 g was required to ensure tag burden (tag mass/fish mass) did not exceed 6% of body weight – a threshold beyond which growth and swimming endurance may be impaired (Table 1;Brown et al., 2006; Collins et al., 2013). Prior to surgery, smolts were anaesthetized in 100-mg/L tricaine methanesulfonate (MS-222) mixed with 200-mg/L NaHCO_3_ buffer until they lost equilibrium and showed no response to overhead stimuli (stage 3 anesthesia; Summerfelt and Smith 1990). Mass (g) and FL (mm) were measured before smolts were transferred into a V-shaped trough on a surgery table. All water baths were aerated, and temperatures were maintained within 2°C of the temperature in the holding tanks at the hatchery. Sterilized transmitters were implanted into a ~5 mm ventral incision that was closed with a single interrupted suture of 4-0 Monocryl (Ethicon, Somerville, NJ, USA). Following tagging, fish recovered until they regained equilibrium and showed a response to overhead stimuli. Tagged fish were held for 24-48 hours prior to transport and release. Water temperature and oxygen were monitored in the transport container using a Polaris C electrochemical oxygen sensor (Oxygaurd International A/S, Denmark). Once at the pumpstation, the fish were introduced to site temperatures by warming holding water at a rate of 0.25°C every 15 minutes through additions of site water. Oxygen levels of 80% or higher were maintained during transport and the warming period. Transmitters were programed with similar transmission rates among years (2024: 45 s; 2025: 35s). No more than 25 acoustic tagged fish were released at a time to avoid code collisions on local receivers. Non-tagged conspecifics (*n =* 20-50) were released with each group to reduce predation pressure on tagged fish. In both years, fish were released downstream of the pumpstation as control fish, or upstream and allowed to pass the through the system independently. In 2025, to isolate the effect of the screw from other components of the infrastructure, fish were released at the top of each pump to pass through the basin and outflow only (Table 3). A group of 45 fish were also released into the Barrowtown Pumpstation over two days in 2025. Dissolved oxygen and water temperature were measured at the time of release using a YSI Pro30 handheld probe. All handling, transport, and surgical procedures complied with Canadian Council on Animal Care (CCAC) guidelines and were approved by the University of British Columbia Animal Care Committee (Protocol # A23-0296).

**Table 1.** Summary of sample sizes (*n*), fork length (FL; mm), and transmitter tag burden (% body mass) for juvenile salmon tagged and released across study years (2024–2025) and release locations. Values represent means with standard deviations [SD] shown in brackets.

| Year | Site | Mean FL [SD]<br>(mm) | Mean Tag Burden (%)<br>[SD] | $n$ |
| --- | --- | --- | --- | --- |
| 2024 | Downstream | 121 [6] | 2.9 [0.5] | 95 |
|  | Upstream | 120 [7] | 2.9 [0.5] | 103 |
| 2025 | Downstream | 122 [5] | 2.9 [0.4] | 114 |
|  | Upstream | 122 [5] | 2.9 [0.4] | 131 |
|  | Outflow_corner | 124 [6] | 2.7 [0.5] | 68 |
|  | Outflow_straight | 122 [5] | 2.8 [0.4] | 71 |
|  | Barrowtown | 126 [7] | 2.9 [0.5] | 45 |

### Array description

Pairs of 69 kHz receivers (VEMCO VR2W; Innovasea Systems Inc, Bedford, Nova Scotia) were installed in the Fraser River. Arrays were deployed in the Mainstem Fraser River at Mission (River kilometer [RKM] 76.8); Barnston Island (RKM 51.3); Derby (RKM 42.8); Port Mann Bridge (RKM 32.5); New Westminster (RKM 26.5) and maintained by Kintama Research, Ltd. Additionally, a single receiver was installed at the outlet of each of pumpstation, and at the confluence of Mountain Slough and the Fraser River. To track subsequent marine migration, detection records from regional acoustic arrays deployed throughout coastal waters – operated by Fisheries and Oceans Canada, the University of Victoria, the Pacific Salmon Foundation, and the Ocean Tracking Network – were retrieved from the Ocean Tracking Network (Figure 1).

### Analysis

#### Data preparation

Time based filtering was applied to the raw acoustic telemetry detections. The first filter removed detections prior to release, and the second removed false positives, identified as closer together than the minimum transmission interval in each year. Finally, any isolated detections on receivers were removed. In total filtering removed 12,350 detections and the final dataset contained 281,365. Modeling was initially attempted with each array fit as a separate location, but the fitted models produced unidentifiable parameters, or negative co-variance matrices. To address this issue, caused by low detection probabilities, we pooled detections between Mission and Port Mann Bridge into a “middle Fraser” array. Detections were also pooled on the marine receivers into a single “marine array”. Including marine detections allowed us to estimate survival to New Westminster. Detection histories were then compiled and arrange chronologically. Individual detection data were structured into spatial capture histories, where each digit represented a receiver array ordered sequentially from upstream release to the marine terminus (1 = detected >= 2 times; 0 = detected <2 times). The final capture history sequence varied among stations as Hammersley had an additional receiver at the confluence of Mountain Slough and the Fraser River (RKM 109; Table 2).

**Table 2.**
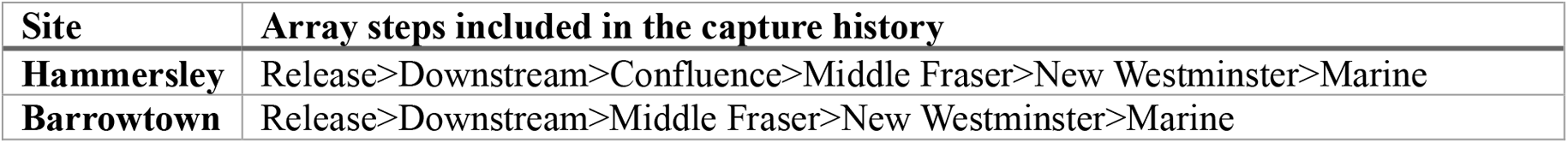
Capture history sequence for fish released at individual pumpstations.

| Site | Array steps included in the capture history |
| --- | --- |
| **Hammersley** | Release>Downstream>Confluence>Middle Fraser>New Westminster>Marine |
| **Barrowtown** | Release>Downstream>Middle Fraser>New Westminster>Marine |

#### Survival analysis

Survival of acoustic-tagged smolts was modeled using a Cormack–Jolly–Seber (CJS) mark-recapture framework for open populations (Cormack 1964; Jolly 1965; Seber 1965). Acoustic detections at successive receiver arrays along the migration route were treated as live recaptures. Reach-specific apparent survival (φ) and array detection probability (*p*) were then estimated using Maximum Likelihood via RMark (Laake J.L., 2013) and Program MARK (White & Burnham, 1999). Assumptions of the CJS model include equal survival and detection probabilities among individuals in a group, and instantaneous sampling approximation (Lebreton et al., 1992). At Hammersley, we expected survival and detection probability may vary among release group and study years. To accommodate heterogeneity, models with φ estimated separately by group and year and *p* estimated by (year, group, and array) were included in the candidate model set (Table 3). Pump operation data, necessary to determine which pump each upstream released fish passed through, was available in 2024 but not 2025. Thus, a combined survival estimate was generated for all fish passing Hammersley pumpstation each year. The survival of fish released into the outflows was modeled separately (straight or corner).

**Table 3.** Cormack–Jolly–Seber (CJS) candidate model selection results evaluating apparent survival (Φ) and detection probability (*p*) for acoustically tagged juvenile salmon migrating past Hammersley and Barrowtown pump stations. Models are ranked by Quasi AIC (QAIC) to account for overdispersion (Hammersley: ĉ= 1.73; Barrowtown: ĉ= 3.80). Model structure, number of parameters (*K*), difference from the top-ranked model (ΔQAIC), Akaike weight (*w*_i_), and model deviance are all also displayed.

| Site | Model Formula | N<br>par | QAIC | ΔQAIC | weight | Deviance |
| --- | --- | --- | --- | --- | --- | --- |
| Hammersley | Phi(~release + array + year)p(~array + year) | 15 | 1238.4 | 0.0 | 1.0 | 72.1 |
|  | Phi(~release + year)p(~array + year) | 11 | 1303.1 | 64.7 | 0.0 | 145.0 |
|  | Phi(~1)p(~array + year) | 7 | 1313.1 | 74.7 | 0.0 | 163.0 |
|  | Phi(~release + array + year)p(~1) | 10 | 1350.9 | 112.4 | 0.0 | 194.7 |
|  | Phi(~1)p(~1) | 2 | 1751.3 | 512.9 | 0.0 | 611.3 |
|  | Phi(~release + year)p(~1) | 6 | 1756.4 | 518.0 | 0.0 | 608.4 |
| Barrowtown | Phi(~1)p(~1) | 2 | 38.2 | 0.0 | 0.9 |  |
|  | Phi(~array)p(~1) | 5 | 42.1 | 4.0 | 0.1 |  |

Barrowtown lacked a matched control group. To evaluate survival at this site, we first parameterized a model with reach scale survival where, φ was estimated separately for the first interval representing pump passage from a pooled downstream Fraser reach. However, this parameter structure resulted in non-positive covariance matrices and unidentifiable parameters due to data sparsity. Next, we developed a candidate model set with data structured by the reaches designated in Table 1. In this set, φ and *p* were allowed to vary by time. However, the model with *p* estimated by time, also resulted in a non-positive covariance matrix due to data sparsity. Consequently, the final model set for Barrowtown was restricted to time varying or pooled φ probabilities and a pooled *p* probability (Table 3). We present reach specific survival estimates for Hammersley and cumulative survival estimates for both stations to New Westminster. The product of reach specific survival was calculated as an estimate of survival to New Westminster, and the variance of these values was estimated using the Delta Method (Equations 5 and 8; Powell 2007).

Goodness-of-fit testing was conducted via parametric bootstrapping. Simulations (n=500) of the fully parameterized model was run for each site to calculate the overdispersion parameter (ĉ). At Hammersley, moderate over dispersion was detected (ĉ= 1.73). At Barrowtown, more severe overdispersion was detected 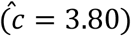. While values of ĉ> 3.0 can indicate unmodeled structure, in the case of Barrowtown it is more likely this value reflects data sparsity (*n* = 45) because all alternative model structures resulted in non-positive covariance matrices or larger values of (ĉ). Candidate models were ranked using QAIC_c_, and all parameter standard errors and confidence intervals were adjusted by 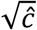 (Burnham & Anderson, 2002).

### Passage and Migration behaviour

Approaches and retreats from the pump forebay were inferred from detections on two receivers installed upstream of the Hammersley pumpstation. One receiver was located in the forebay with the hydrophone directed at the entrance to the pumpstation and a second receiver was placed 30 meters upstream of the pumpstation also directed toward the pump intake (Figure S1). Discrete approach events were identified by a gap of 60 minutes or more between detections on either of these receivers. We then summarized the median and interquartile range of the number of approaches made by coho salmon successful or unsuccessful in passing the pumpstation. We also summarized time between approaches to median and IQR. The approach summaries were generated from the 2025 data only because only one fish failed to pass in 2024.

Passage delays and migration behaviour were quantified using the receivers downstream of the pumpstations. We estimate pump related migration delays, from fish released upstream and detected on receivers immediately downstream of the stations. For this summary, we calculated the difference between release and the first detection downstream. We estimated migration speed (kilometers/day) for each reach, and total time to New Westminster for each group in each year. Fish migrating from Hammersley traveled 87.3 km to New Westminster, while fish leaving Barrowtown migrated 60.3 km. The time in days to reach New Westminster was divided by the distance each group had to travel to arrive at kilometers/day. The District of Kent provided pump operation data for the Hammersley station in 2024 which was summarized to the number of hours per day at least one pump operated. Similar data was not available in 2025 due to a computer outage.

## Results

### Environmental Conditions

Discharge in the Fraser River varied among years. In 2024, freshet was late arriving and slower to progress resulting in discharges between 2800 m^3^s^−1^ and 3500 m^3^s^−1^ for the period this study operated. In 2025, discharge was higher (3500-3800 m^3^s^−1^) and matched historic freshet conditions (Figure S3). Water temperatures ranged from 13°C to 17°C among release locations and years (Figure 2). In 2024, upstream and downstream release groups experienced similar environmental conditions. In 2025, fish released upstream at Hammersley experienced low dissolved oxygen (< 50 %) compared to those released downstream or at Barrowtown.

**Figure 2.**
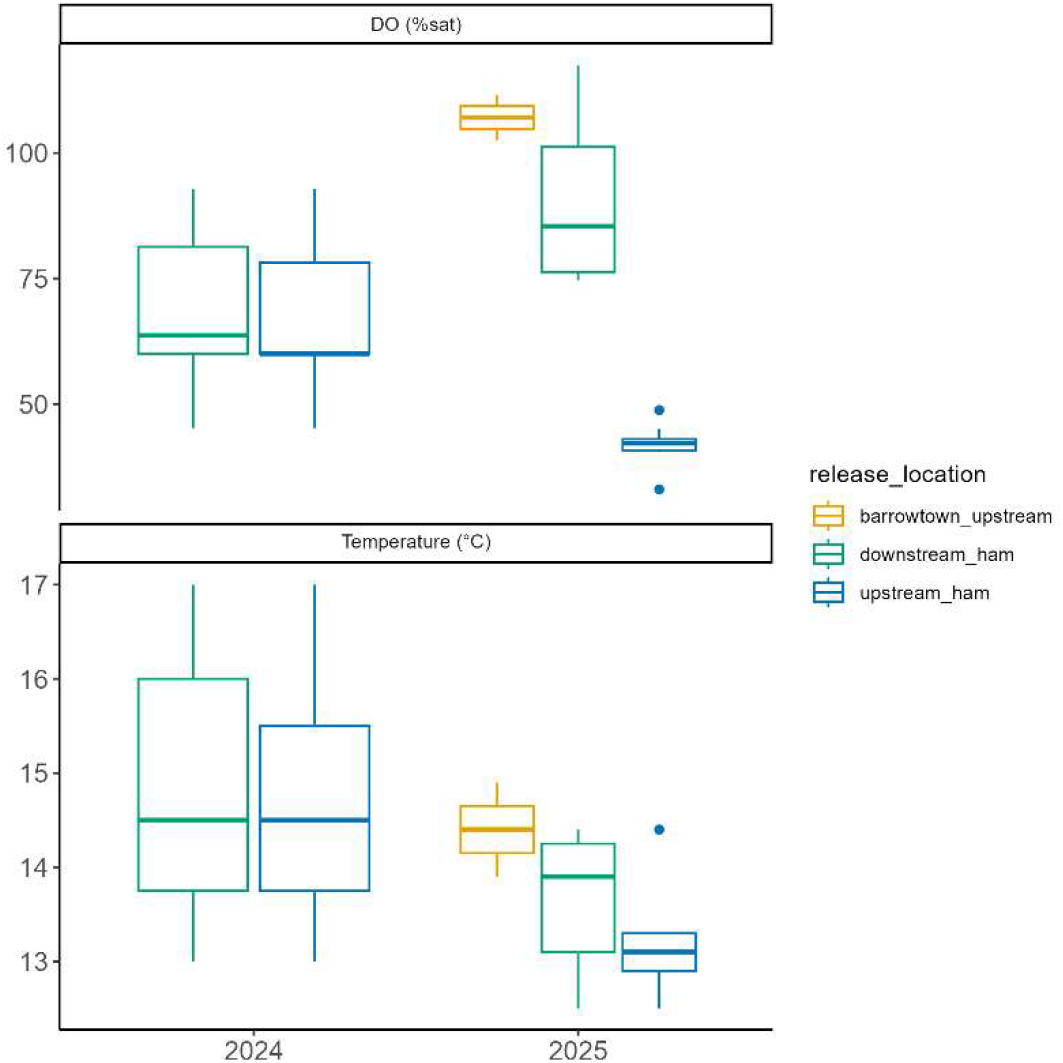
Dissolved oxygen and water temperatures at the time of release for acoustically tagged yearling coho salmon. Releases occurred at the Hammersley open Archimedes pumpstation (X_ham; Agassiz, British Columbia) or the Barrowtown pumpstation (Abbotsford, British Columbia).

### Survival estimates

For Hammersley, the best ranked model included φ estimated by release location, year and array and *p* estimated by array and year. The closest ranked model had a ΔQAIC of 64.7, indicating no support for alternative model structures. The cumulative survival from release to New Westminster for downstream released fish was similar among years (2024:39.3% [95% CI: 32.7-46.4]; 2025:41.6% [95% CI:34.5-49.1]; Figure 2). Upstream released fish also had similar, but lower survival (2024:27.0% [95% CI: 21.6-33.9]; 2025:29.9% [95% CI:23.5-35.9]) than downstream released fish in both years. Fish passing just the outflows experienced a slightly slower survival than fish passing the entire pumpstation (3-6% reduction, Figure 3). Among groups released at Hammersley, the largest differences in survival materialized in reach 3, between the confluence of Mountain Slough and the Middle Fraser River (Figure 4). The Null model, with pooled survival and detection probability, was the best ranked model for fish passing Barrowtown. Cumulative overall survival for Barrowtown pumpstation was 12.5% (95% CI: 0.0-0.4; Figure 3). For fish released at Barrowtown, the reach specific survival was estimated to be 50.3% (95% CI:29.1-71.3; Figure 4).

**Figure 3.**
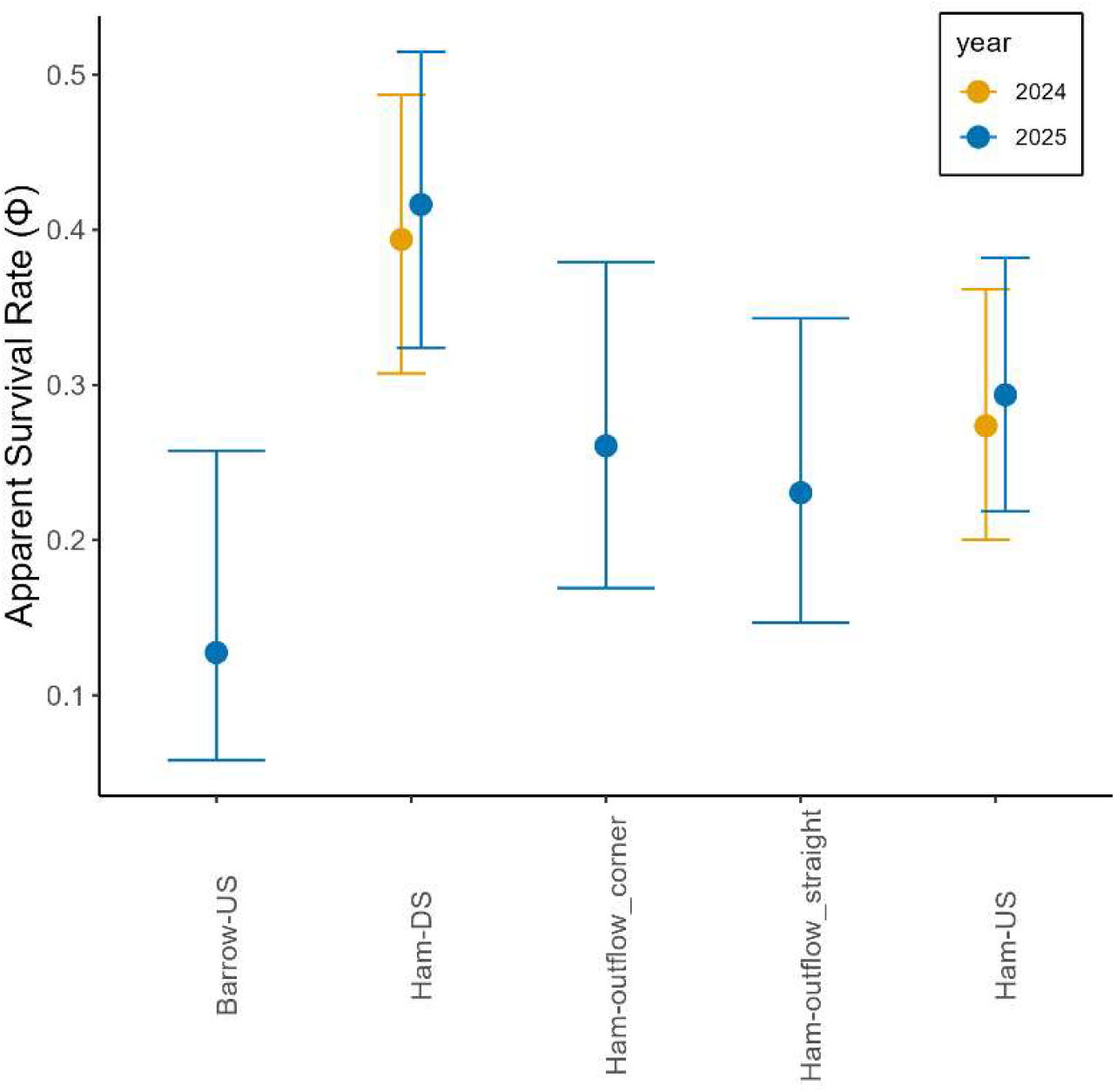
Cumulative apparent survival between release location and the Fraser River at New Westminster. Ham = Hammersley open Archimedes screw pump in Agassiz, British Columbia (DS= downstream [control]; US = Upstream; Outflow_X=individual outflows of each of the paired pumps). Barrowtown refers to Barrowtown submersible pumpstation in Abbotsford, British Columbia.

**Figure 4.**
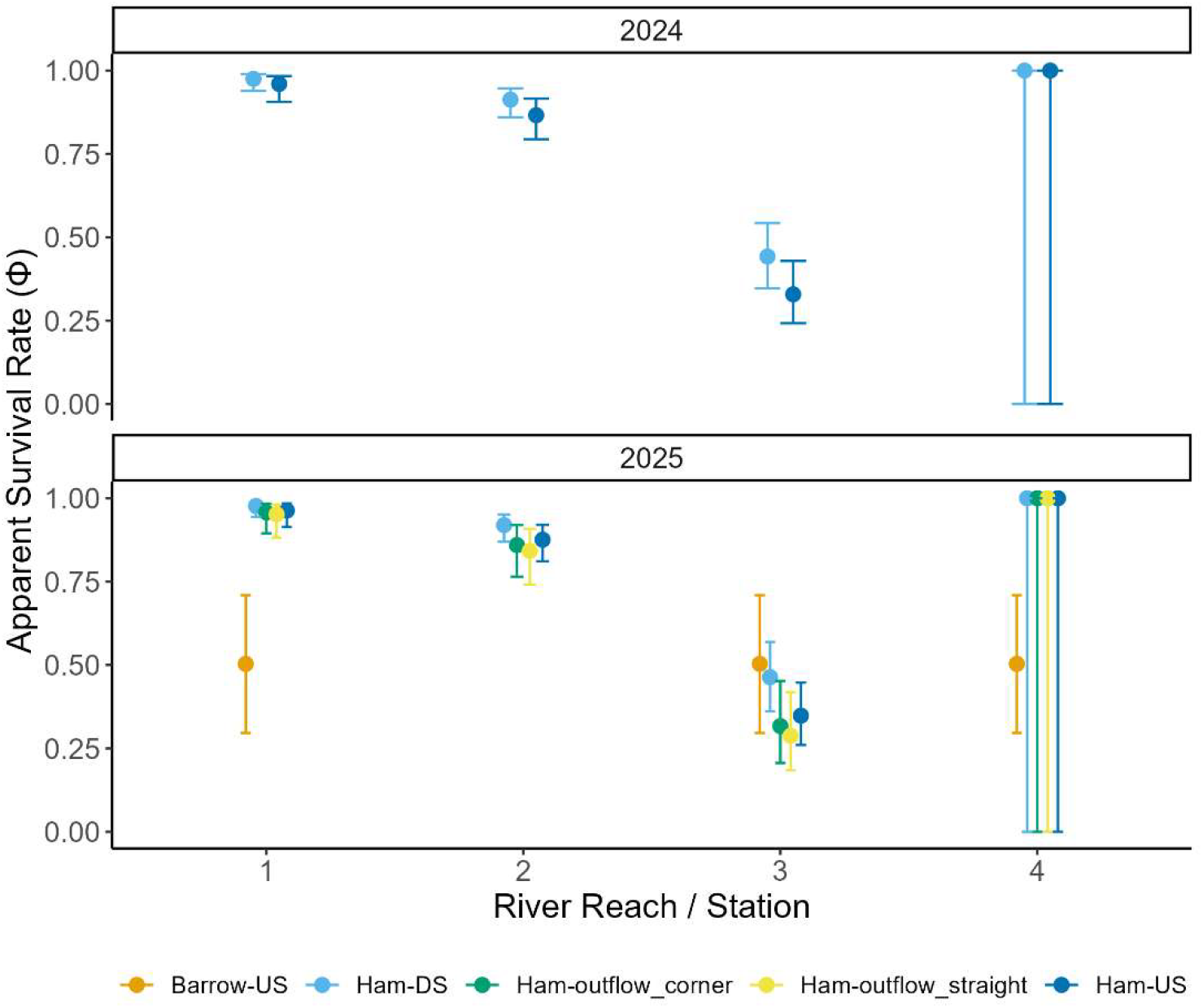
Reach scale apparent survival between release location and the Fraser River at New Westminster. Ham = Hammersley open Archimedes screw pump in Agassiz, British Columbia (DS= downstream [control]; US = Upstream; Outflow_X=individual outflows of each of the paired pumps). Barrowtown refers to Barrowtown submersible pumpstation in Abbotsford, British Columbia.

#### Detection Probability

Detection probability for fish released at Hammersley was lowest at the New Westminster array in both years (2024: 26.8% [95%CI:18.6-37.1]; 2025: 8.2% [95% CI:5.2-12.9]; Table 4). In 2024, detection probability ranged from 92.2%-99.8% for the receivers upstream of New Westminster. In 2025, these stations were estimated to have detection probabilities between 73.8 and 99.1% (Table 4). The pooled detection probability across all arrays for fish released at Barrowtown was 89.8% (95% CI:32.4-99.3). The utility of including coastal marine arrays was particularly evident for this group: while only a single individual was registered at the New Westminster array, four fish were subsequently detected across the marine network, confirming out-migration past the river terminus.

**Table 4.** Detection probability estimates (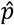, %) and 95% confidence intervals across acoustic receiver arrays at in the Fraser River between either Hammersley or Barrowtown pumpstation in 2024 and 2025.

| Site | Year | Array | Estimate | 95% LCL | 95% UCL |
| --- | --- | --- | --- | --- | --- |
| Hammersley | 2024 | Downstream pump | 92.0 | 88.0 | 94.7 |
|  |  | Confluence | 99.8 | 98.4 | 100.0 |
|  |  | Middle Fraser | 96.7 | 92.2 | 98.7 |
|  |  | New Westminster | 26.8 | 18.6 | 37.1 |
|  | 2025 | Downstream pump | 73.8 | 68.8 | 78.2 |
|  |  | Confluence | 99.1 | 94.1 | 99.9 |
|  |  | Middle Fraser | 87.9 | 75.9 | 94.4 |
|  |  | New Westminster | 8.2 | 5.2 | 12.9 |
| Barrowtown | 2025 | Pooled | 89.8 | 32.4 | 99.3 |

### Migration behaviour

Coho salmon successful in passing Hammersley pumpstation approached the pump fewer times (*n*=116; median=8; IQR=4-18) than fish that were unsuccessful (*n*=15; median=16; IQR=9-28). The median time interval between approach events was similar among successful (median=5.2 hours; IQR=3.4-8.2) and unsuccessful (median=5.4 hours; IQR: 3.7-13.8) fish. Fish released upstream of Hammersley took twice as long to pass the pumpstation in 2024 (mean=7.0 days; sd=5.6; *n*=56) as in 2025 (Mean=3.5 days; SD=3.88; n=57). At Barrowtown, passage time averaged 2.5 days (sd=4.65; *n*=21). Migration time between the Hammersley pumpstation (rkm 110.2) and New Westminster (rkm 26.5) varied among years. In 2024, fish released upstream took twice as long to reach New Westminster (Table 5) as fish released downstream and migration speed was higher across all but one reach for downstream fish (Figure 5). In 2025, among group differences in travel speed were smaller (Figure 5). None of the fish released into the outlet with a corner were detected at New Westminster. The single fish from Barrowtown detected on the New Westminster array took 4 days to reach this location (Table 5). In May of 2024, the Hammersley pumpstation operated 1.0 to 7.0 hours per day (Figure 6). Peak operations occurred in June after all study fish had left the area (Figure 6).

**Table 5.** Time (days) for fish to reach New West Minster from their release location. Fish released at Hammersley Archimedes Pumpstation (Ham) migrated 83.7 km while fish released at Barrowtown Pumpstation (Barrow) migrated 60.3 km.

| year | Release location | Mean days reach New Westminster [sd] | Speed (km/day [sd]) | *n* |
| --- | --- | --- | --- | --- |
| 2024 | Ham-DS | 9.9 [1.3] | 66.0 [8.4] | 4 |
|  | Ham-US | 19.4 [6.7] | 12.5 [4.3] | 7 |
| 2025 | Barrow-US | 5.7 [-] | 10.7 [-] | 1 |
|  | Ham-DS | 9.7 [6.3] | 13.4 [8.7] | 6 |
|  | Ham-US | 10.3 [4.4] | 19.0 [8.1] | 6 |
|  | Ham-outflow_straight | 5.0 [2.6] | 32.4 [16.6] | 4 |

**Figure 5.**
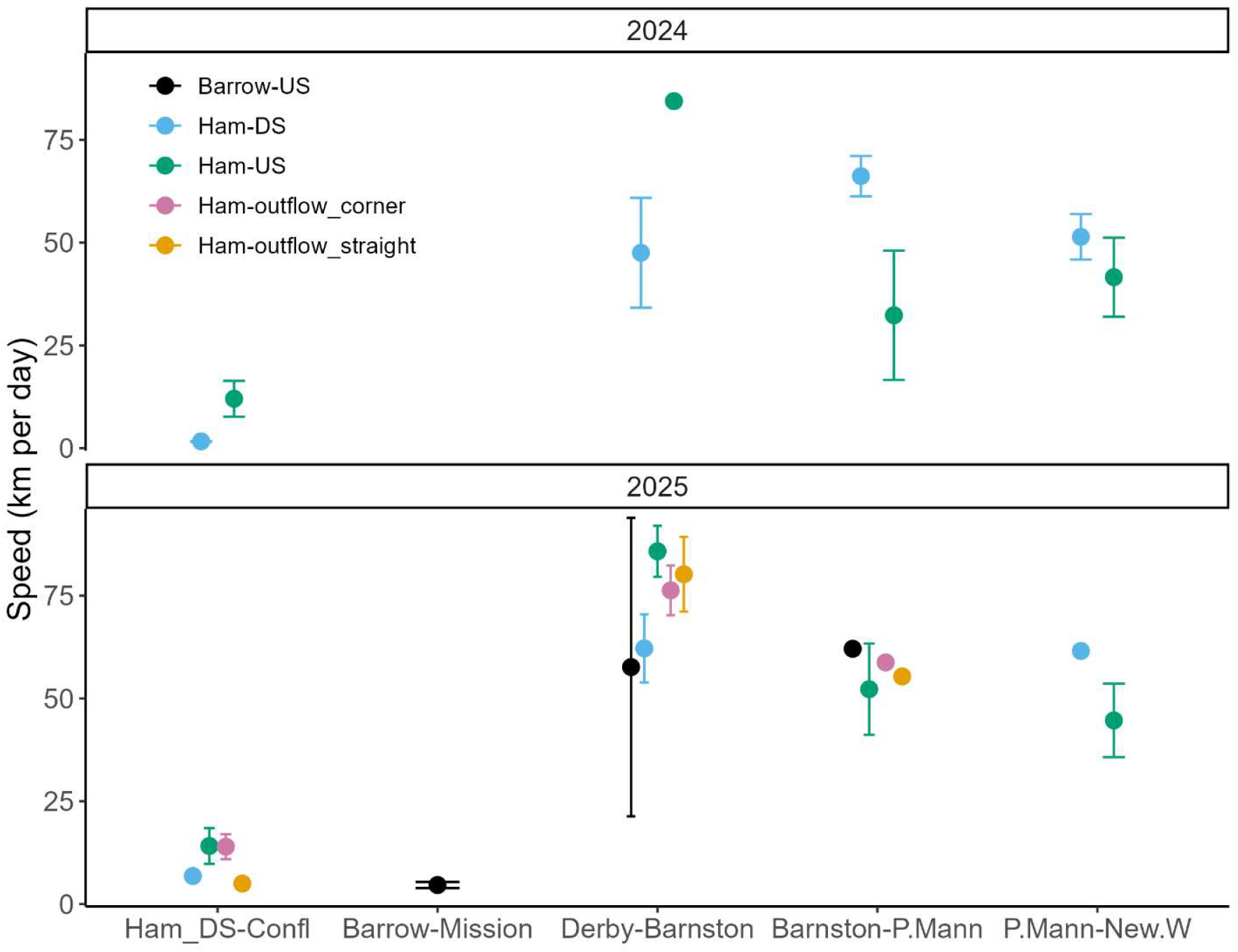
Mean travel speed (km/day) per reach for juvenile coho salmon in the Fraser River. Release groups above (Barrow-US, Ham-US) and below (Ham-DS) flood pumpstations. Ham= Hammersley open Archimedes pumpstation, Barrow= Barrowtown radial pumpstation. Ham-outflow_corner & Ham-outflow_straight refer groups released into the outflow of individual screw pumps at Hammersley. Points indicate mean, and error bars represent standard errors. The absence of an error bar indicates a sample size of one for the particular group and reach.

**Figure 6.**
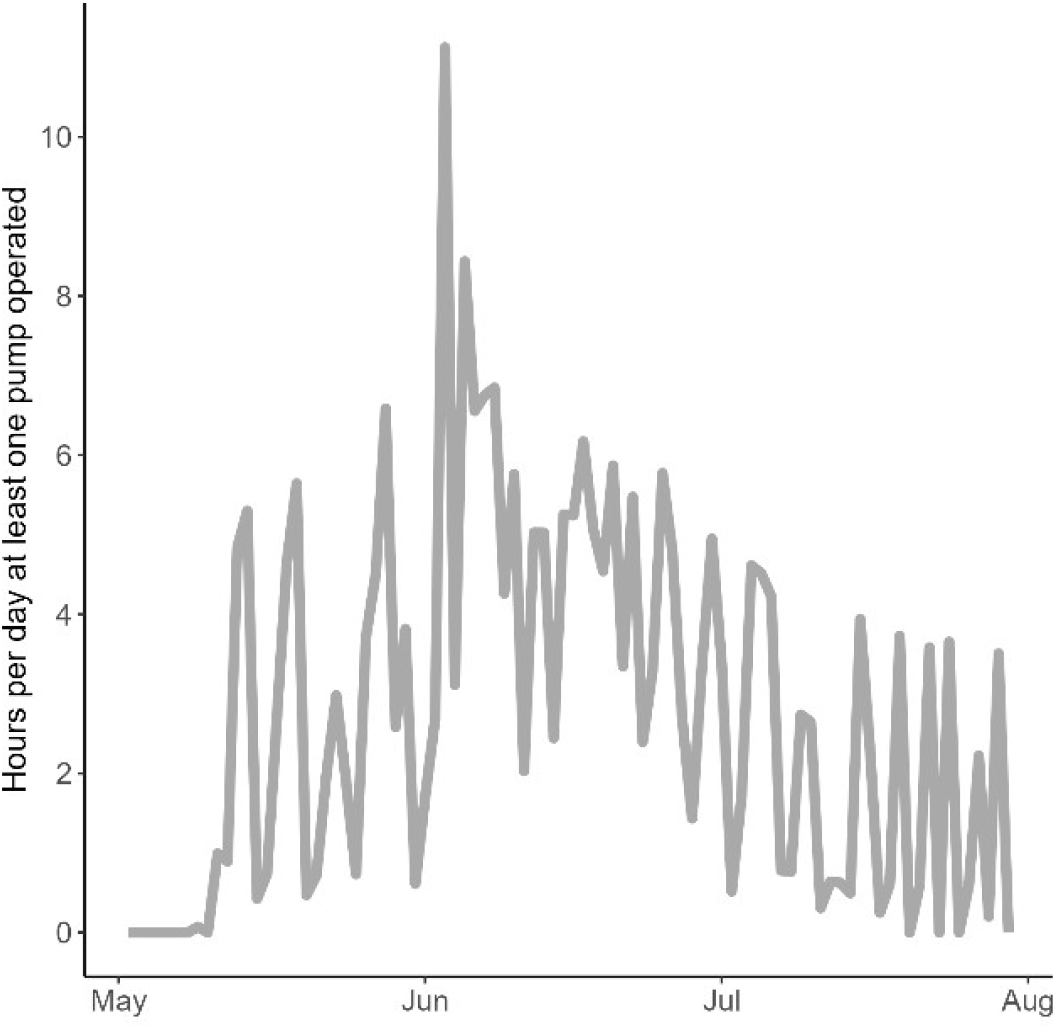
Daily operations of the Hammersley open Archimedes screw pumpstation in Agassiz, British Columbia. The station contains two screw pumps. Data is summarized to represent the number of hours per day at least one pump was operated in spring and summer of 2024.

## Discussion

Juvenile coho salmon survival was compromised by entrainment through both the open-Archimedes and conventional radial pumps. Baseline natural mortality for juvenile coho salmon migrating between Mountain Slough (rkm 110) and New Westminster (rkm 26.5) averaged 60% across years. In contrast, coho entrained through the conventional radial station at Barrowtown experience 87.5% cumulative mortality between the Barrowtown (rkm 87.4) station and New Westminster. Mortality directly attributable to the open-Archimedes pumpstation averaged 11.5% across years, representing a 15% improvement over the conventional station. Counterintuitively, groups released directly into the outflow of the open-Archimedes station experienced a small (2-3%) reduction in survival compared to fish passing the entire station. This discrepancy may reflect the compounding stress of transport and exposure to a hydraulic challenge, as outflow fish were released directly to the pump with no recovery time following transport. While the open Archimedes pump improved survival over the conventional radial pump, substantial mortality (> 10%) is associated stations employing this technology. Survival impacts due to pump entrainment were not immediate; all groups released at the open-Archimedes station exhibited high survival from release to the receivers downstream the station (95-98%). Survival differences manifested as fish entered the Fraser River, both the full-station and outflow groups experienced ~5% declines in survival relative to controls. The divergence in survival peaked between the confluence and Middle-Fraser (Mission to Port Mann Bridge) where survival differed by 16% between control fish and those exposed to the full-station or outflows. If coho juveniles survived to the Middle-Fraser, reach specific survival stabilized to 99% to across all groups. The delayed divergence in survival for fish passing the open-Archimedes pump indicates the biological costs of passage may be predominantly latent, manifesting only when compromised fish encounter selection pressures.

Entrainment through pumps also affected migration timing and travel speeds, which likely contributed to reach-specific survival disparities. In 2024, coho juveniles released upstream of the open-Archimedes pumpstation took 10 days longer to reach New Westminster than downstream controls; however, this discrepancy narrowed to one day in 2025. Coho released directly into the station outflows exhibited the most rapid transit overall, reaching New Westminster in 5 days—twice as fast as both the control and full-station passage groups. The prolonged migration times observed for upstream fish in 2024, resulted from substantial passage delays and slower reach specific travel speeds. In contrast, 2025 migration speeds are largely uniform across groups and reaches, with the exception of the Derby to Barnston reach where upstream fish exhibited accelerated transit time. The single reach spike in velocity compensated for the 3.5-day delay in passing the station. It is impossible to distinguish the mechanism responsible for the much faster pump passage times in 2025 as operations data were not available; however, it is likely the pump operated more frequently in in this year due to a greater low elevation snowpack and more typical early spring precipitation pattens.

Post-passage mortality for juvenile coho salmon entrained through an open Archimedes screw was substantially higher – by a factor of three to ten – than rates previously documented for closed-fixed designs. In the only prior evaluation of an Archimedes pump with Pacific salmon, McNabb et al. (2003) reported just 1% mortality for sub-yearling Chinook Salmon at an experimental pumpstation in Red Bluff, California. However, that evaluation routed juvenile Chinook Salmon directly into holding boxes to quantify survival over a 96-hour holding duration. While holding studies capture acute physical trauma, latent mortality often does not manifest until days or weeks after passage, when secondary ecological stressors expose the performance costs of sub-lethal injury (Burnett et al., 2017). Furthermore, sub-yearling salmon (< 74 mm), which may be more resilient to hydraulic forces, were the focus of the Red Bluff tests (Cox et al., 2023). Beyond methodological differences, the pumpstations differed in physical design, the largest being the type of screw pump installed. To our knowledge our study is the first to test an open-Archimedes pump with Pacific salmon and the mortality rates observed here align those for other physostomous species (*Abramis brama* and *Rutilus rutilus*) at another open-screw station (Pauwels et al., 2020). Due to the complexity of pumpstation designs and range of features that have the potential to cause harm to fish, the results presented here are not directly transferable to other stations using open-screw designs. Further research on the sensitivity of individual species to various hydraulic forces is required to guide the design of fish-safe drainage infrastructure.

This study provides a rare empirical estimate of baseline natural mortality for hatchery origin coho salmon yearlings in the lower Fraser River. Across both study years, natural mortality was high, averaging 60% with in an 84 km reach. By comparison, wild juvenile sockeye salmon (O. nerka) migrating from Chilco Lake to the mouth of the Fraser River exhibited substantially lower mortality (~40%) over a distance seven times longer than the reach evaluated here (Stevenson et al., 2019). We employed standardized surgical procedures and strict transport guidelines to minimize stress. Based on empirical holding studies using the identical surgical protocols, handling and tag burden likely account for 10-15% of total observed losses in this study (Collins et al., 2013b; Stevenson et al., 2019). Consequently, the observed baseline mortality likely reflects the acute ecological hazards of the lower Fraser River. This is a heavily modified reach presented migrating smolts with an array of compounding stressors including high predator densities (Seifert & Moore, 2018); contaminants (Lo et al., 2026); and widespread habitat loss (Chalifour et al., 2022; Finn et al., 2021). That yearling coho salmon experience high rates of natural mortality through this reach is an important finding towards protecting vulnerable stocks migrating through this region.

### Study Limitations

Several logistical and environmental constraints warrant consideration when interpreting these findings. First, receiver detection efficiency at our downstream boundary array (New Westminster) varied substantially between years, declining from 26.8% during low discharge in 2024 to 8.2% at typical freshet discharge. While the multi-state Cormack–Jolly–Seber framework mathematically accounts for imperfect detection by leveraging downstream marine arrays, these diminished detection probabilities inevitably broadened confidence intervals on terminal reach survival estimates. Second, sample sizes at the conventional Barrowtown pumpstation were modest (n= 45) resulting in elevated model overdispersion which can bias standard errors; however, we accounted for this through the use of QAIC. Despite these limitations, the observed survival estimates serve as important benchmarks for a little studied type of infrastructure.

#### Management & Engineering Implications

Modernizing aging drainage infrastructure with “fish-friendly” technology is frequently championed as beneficial for both flood protection and salmon restoration, yet our results demonstrate that replacing conventional pumps with open-Archimedes screws does not eliminate ecological hazards. While the open-Archimedes design conferred a 15% survival benefit over legacy radial pumps, a mortality penalty of 11.5% remains unsustainable in floodplains where out-migrating smolts already face high baseline losses (~60%). Because mortality manifested primarily as delayed, latent losses in the mainstem river – likely driven by sublethal trauma from shear stress, turbulence, and structural collisions – impact assessments relying solely on acute holding trials will systematically underestimate population-level effects. To mitigate both acute and latent losses, regional diking districts and fisheries managers must transition from relying exclusively on installation of “fish-friendly” pumps toward holistic hydraulic management that prioritizes timely passage, total facility safety (including tailrace hydraulics), and non-entrainment bypass routes whenever possible.

## Acknowledgements

This project was financially supported through the BC Salmon Restoration and Innovation Fund, the National Science and Engineering} Council (NSERC) of Canada and mitacs. Field and logistical support was provided by the Cheam First Nation, Sumas First Nation, Resilient Waters, Watershed Watch Salmon Society, Bailey Environmental, Talik Ecological, and Chehalis River Hatchery. The authors are grateful for the opportunity to work in the traditional territories of both the Cheam First Nation and Sumas First Nation. The project would not have been possible with out the support of David Welch and Kintama Research.

