## Supplemental Figures S1_S3 for "Passage through submersible and Archimedes pumps results in decreased migration survival in coho salmon yearlings"

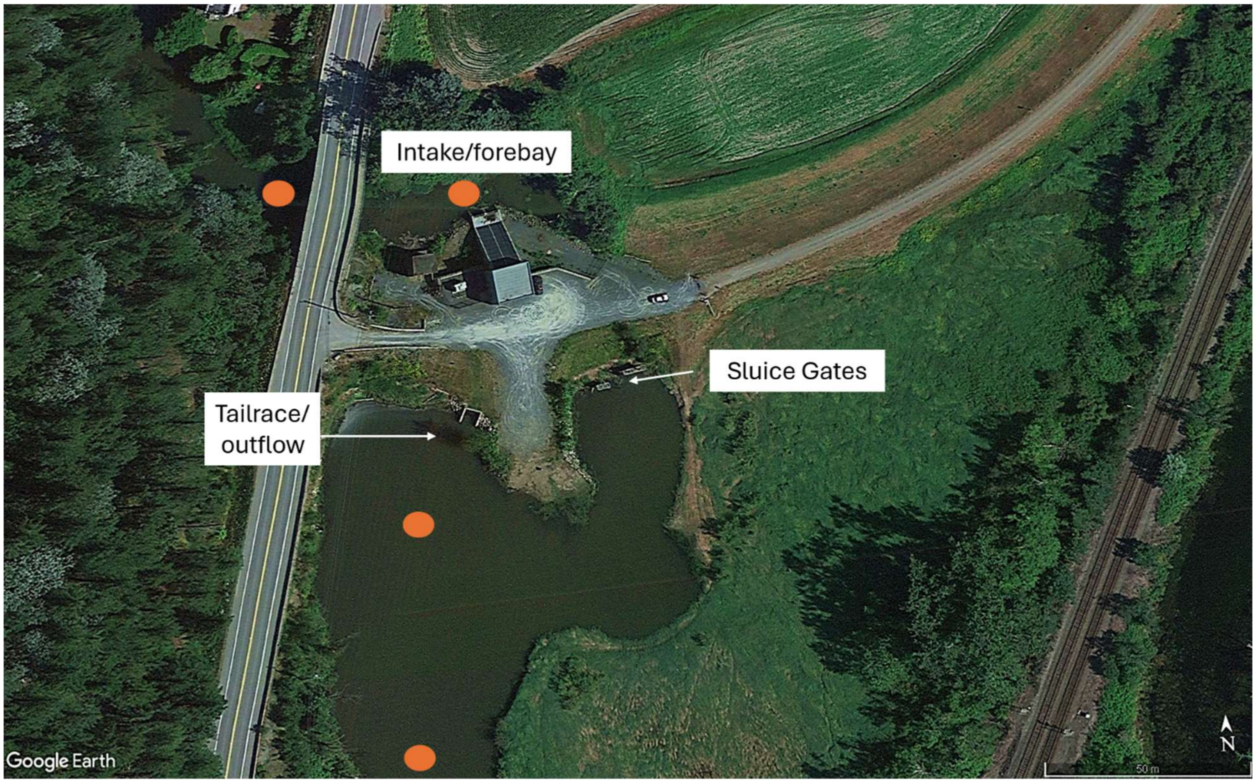


Figure S1 Hammersley open Archimedes screw pumpstation from above. Orange circles indicate acoustic receivers placed around the station.

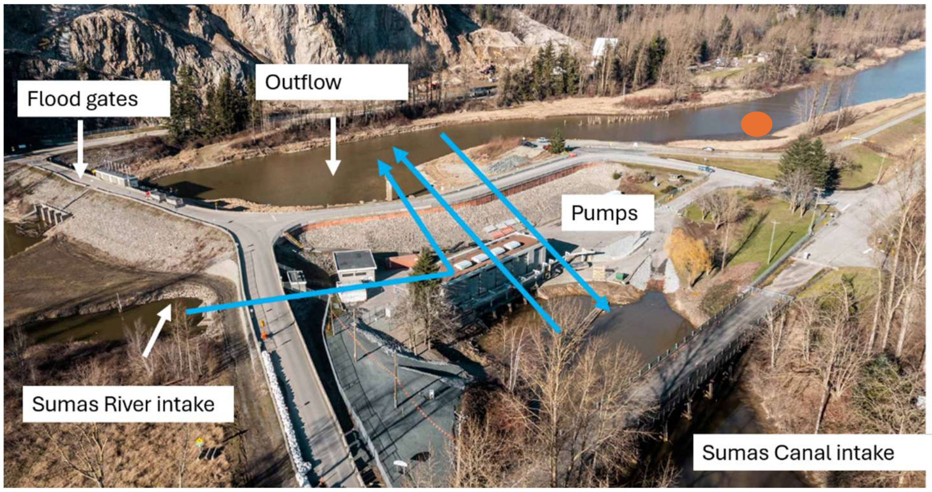


Figure S2. Barrowtown conventional radio pumpstation. Blue arrows indicate direction water moves through the station. Orange circle indicates location of receiver placed downstream to detect passage events. Coho salmon were released into the Sumas River intake in May 2025.

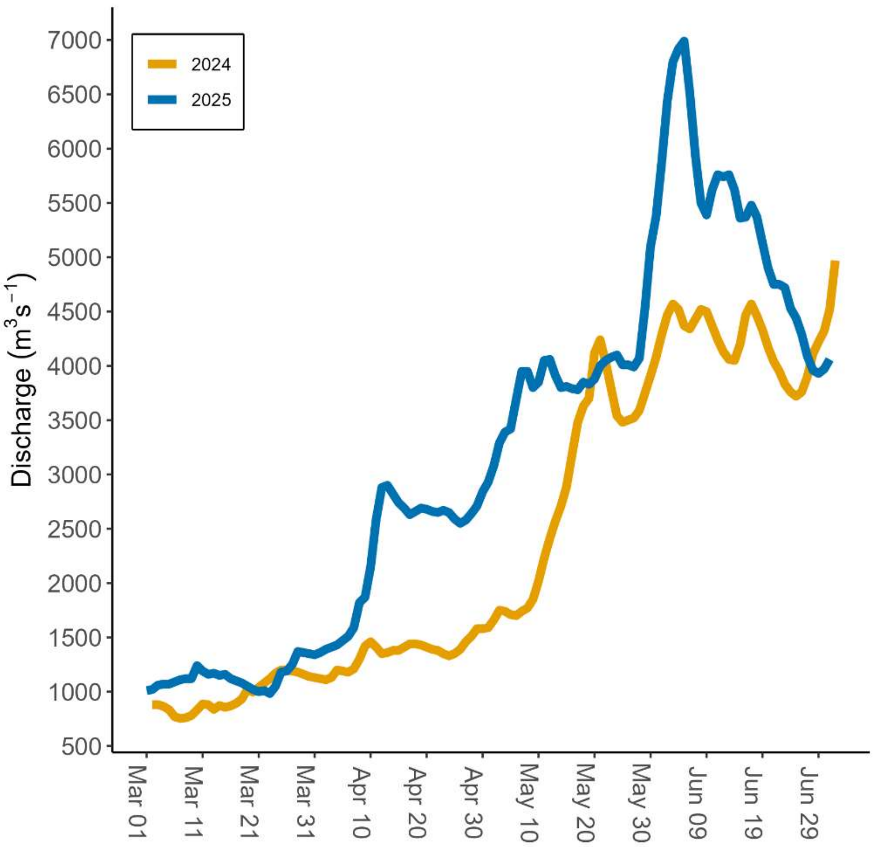


Figure S3. Average daily discharge for the Fraser River at Hope (Water Service of Canada Gauge: 08MF005) in the spring and summer of 2024 and 2025.
